# Enterovirus Evolution Reveals the Mechanism of an RNA-Targeted Antiviral and Determinants of Viral Replication

**DOI:** 10.1101/2023.02.20.529064

**Authors:** Jesse Davila-Calderon, Mei-Ling Li, Srinivasa R. Penumutchu, Christina Haddad, Linzy Malcolm, Amanda E. Hargrove, Gary Brewer, Blanton S. Tolbert

## Abstract

Selective pressures on positive-strand RNA viruses provide opportunities to establish target site specificity and mechanisms of action of antivirals. Here, Enterovirus-A71 revertant viruses with resistant mutations in the SLII IRES domain (SLII^resist^) were selected at low doses of the antiviral DMA-135. The EV-A71 revertant viruses were resistant to DMA-135 at concentrations that robustly inhibit replication of wild-type virus. EV-A71 IRES structures harboring the suppressor mutations induced efficient expression of reporter Luciferase mRNA in the presence of non-cytotoxic doses of DMA-135 whereas DMA-135 dose-dependently inhibited Luciferase expression from the wild-type IRES element. NMR studies indicate that the resistant mutations change the structure of SLII at the bulge loop binding site of DMA-135 and at part of an extended surface recognized by host RNA-binding protein AUF1. Comparisons of biophysical analysis of complexes formed between AUF1, DMA-135, or either SLII or SLII^resist^ show that DMA-135 stabilizes a ternary complex with AUF1-SLII but not AUF1-SLII^resist^. Further studies demonstrate that the hnRNP A1 protein retains binding affinity for SLII^resist^, illustrating that DMA-135 inhibition and viral resistance do not perturb the SLII-hnRNP A1 arm of the regulatory axis. Taken together, this work demonstrates how viral evolution under selective pressures of small molecules can elucidate RNA binding site specificity, mechanisms of action, and provide additional insights into the viral pathways inhibited by the antiviral DMA-135.

## Introduction

Non-polio human enteroviruses (EV) are positive strand RNA pathogens that pose a serious threat to global economies and healthcare infrastructures. EVs infect millions of people around the world and thousands in the United States annually. Although infections typically manifest with mild and self-limiting illness, prolonged infections in the immunocompromised can lead to severe neurological disorders, cardiopulmonary failure, and death.(*1–7*) Thus, EV infections have the potential to develop into severe health outcomes given that disease pathogenesis impairs multiple organ systems.(*8*)

Transmission of EV-A71, an etiological agent of the Hand, Foot, and Mouth Disease (HFMD), has become endemic to the Asia-Pacific region with major outbreaks every 3-4 years such as a 2018 epidemic in Vietnam where more than 53,000 hospitalizations and six deaths were reported.(*9*) The National Institute of Allergy and Infectious Diseases (NIAID) recognized EV-A71 and related EV-D68 as group II re-emerging pathogens. As of the time of writing, treatment of EV-A71/D68 infections remains largely supportive because there are no FDA-approved vaccines or therapeutics, emphasizing the need to develop a comprehensive understanding of the molecular mechanisms involved in host-virus interactions.

EV-A71 is a non-enveloped, single-stranded, positive-sense RNA virus found in species A of the *Enterovirus* genus within the *Picornaviridae* family. Its 7,500-nucleotide genome serves as template for viral translation and replication by partitioning these functions via changes to the host-virus interactions that regulate the cellular stages of viral replication.(*10*) The viral genome encodes a single 250-kDa polyprotein using a long open reading frame (ORF) that is flanked by highly structured untranslated regions (UTRs). Given this limited coding capacity, EV-A71 coordinates complex molecular events to usurp host proteins to drive translation of the viral polyprotein.(*11*) Notably, EV-A71 utilizes a type-I internal ribosome entry site (IRES) within its 5’UTR to initiate translation in a cap-independent mechanism assisted by cellular RNA-binding proteins (RBPs), collectively known as ITAFs (IRES Trans Acting Factors). (*12, 13*) IRES mediated translation commences immediately following infection where ITAFs modulate the efficiency by which the ribosome loads internally onto the 5’UTR.(*14*) Thus, IRES-ITAF interactions are essential determinants of the earliest fates of EV-A71 replication, making them attractive targets for therapeutic intervention.(*11, 15*)

The IRES of EV-A71 is predicted to fold into five major stem loops (SL II-VI) in addition to the 5’-end cloverleaf structure required for virus replication. SLII is the only domain whose structure is solved at high resolution, and several of its binding partners functionally validated.(*15–21*) The structure of EV-A71 SLII consists of a phylogenetically conserved 5-nt bulge and 7-nt apical loop.(*16, 21*) Mutations or deletions to the bulge sequence impair viral replication by attenuating IRES-dependent translation.(*16, 21*) SLII binds several cellular RNA-binding proteins (RBPs) and a viral-derived small RNA vsRNA1.(*16–21*) Of the RBPs, hnRNP A1 and AUF1 are essential ITAFs that exert regulation by competitively binding SLII to differentially modulate viral translation levels.(*19, 20*) HnRNP A1 stimulates IRES-dependent translation, whereas AUF1 antagonizes binding of hnRNP A1 to downregulate polyprotein synthesis. In a recent study, we leveraged the functional significance of the SLII IRES domain to screen a library of small molecules and identified the dimethylamiloride DMA-135 as a potent inhibitor of EV-A71 replication.(*15*) We demonstrated that DMA-135 attenuates IRES-dependent translation by binding the bulge loop to allosterically stabilize a (DMA-135)-SLII-AUF1 ternary complex that we posited disrupts the homeostatic balance of the SLII-host regulatory axis.(*15*)

Herein, we harnessed the power of viral evolution to establish the cellular mechanism of DMA-135 and to reveal new insights into EV biology. By treating EV-A71 infected cells with an inhibitory dosage of DMA-135, we were able to select for viruses that grow to high titers after 10 rounds of serial passage. Sequencing the 5’UTR of a plaque-purified virus revealed that the revertant mutations mapped to sites adjacent to the bulge motif in the SLII domain, the binding site of DMA-135. (*15*) Specifically, residues C132 and A133 were changed to G132 and C133 in a non-compensatory manner in the resistant SLII RNA (SLII^resist^). Genetically engineered EV-A71 harboring only the C132G and A133C mutations were shown to replicate with uncompromised efficiencies even when exposed to DMA-135 concentrations that completely inhibit the wild-type virus. Dual Luciferase reporter constructs that contain SLII^resist^ retained normal IRES-dependent Luciferase activities with and without inhibitory levels of DMA-135.

To better understand the functional specificity of DMA-135, we carried out a comparative biophysical analysis of wild-type SLII and the resistant mutant. NMR studies indicate that the resistant mutations change the structure of SLII proximal to the bulge loop while preserving the overall folding arrangement of the upper helix, including the 7-nt apical loop. Calorimetric titrations determined that hnRNP A1 binds SLII^resist^ with comparable thermodynamic properties to the wild-type stem loop. By contrast, AUF1 does not bind the resistant mutant detectably by calorimetry or by a biochemical pull-down assay. As anticipated, DMA-135 does not promote the formation of an allosteric ternary complex with AUF1 and the SLII^resist^ construct. Collectively, these results support that the mechanism of action of DMA-135 is to tip the SLII-host regulatory axis towards significantly lower levels of IRES-dependent translation, and the virus can compensate by evolving mutations that restore homeostasis.

We also show that DMA-135 can inhibit replication of the related EV-D68 Fermon variant (*22*) albeit less efficiently than EV-A71. NMR comparison of the SLII structures of EV-A71 and EV-D68 reveal the RNAs adopt similar global folds but different structures within the vicinity of the DMA-135 binding epitope. Notably, EV-D68 consists of an internal loop in place of the 5-nt bulge found in EV-A71. Not only is the internal loop topologically different than the bulge, but its sequence composition varies from the high affinity AUF1-binding motif found in SLII from EV-A71. When taken together, the work here defines the antiviral mechanism of action of DMA-135; it demonstrates that functional specificity can be modulated through natural and drug-dependent viral evolution; and it shows how small molecules can reveal new insights about host-virus interfaces that regulate early stages of EV replication. We expect such studies will prove beneficial for efforts to target viral RNA structures or complexes, and to develop chemical biology reagents that can inform on the cellular stages of viral replication.

## Materials and Methods

### RNA preparation

All SLII constructs used in this study were prepared by *in vitro* transcription using recombinant T7 polymerase that was overexpressed and purified from BL21 (DE3) cells. Synthetic DNA templates corresponding to the EV-A71 2231 and EVD68 isolates, or mutant constructs were purchased from Integrated DNA Technologies (Coralville, IA). Transcription reactions were performed using standard procedures and consisted of 3-6 mL reaction volumes containing unlabeled rNTPs or (C^13^/N^15^)-labeled rNTPs. Following synthesis, samples were purified to homogeneity by denaturing polyacrylamide gel electrophoresis (PAGE), excised from the gel, electroeluted, and desalted via exhaustive washing of the samples with RNAse-free water using a Millipore Amicon Ultra-4 centrifugal device. Samples were annealed by heating at 95 °C for 2 minutes and flash-cooled on ice. Samples were subsequently concentrated and exchanged into 10 mM K2HPO4 (pH 6.5) and 20 mM KCl, 4mM TCEP, and 0.5 mM EDTA using a Millipore Amicon Ultra-4 centrifugal filter device. The concentration of the samples was determined using the respective RNA theoretical molar extinction coefficient, and NMR samples ranged from 0.1 – 0.2 mM at 200 μL.

### Protein Purification

The AUF1-RRM1,2 (residues 70-239 correspond to the p37 isoform) protein construct used in this study was subcloned into a pMCSG7 vector, and subsequently overexpressed as an N-terminal (His)6-tagged fusion protein in BL21 (DE3). Cells were grown to an OD600 of ∼1.0 at 37 °C and induced with 0.5 mM IPTG. Immediately after induction, cells were cooled to 20 °C and harvested by centrifugation 18 hours post-induction. The respective (His)6-tagged A-RRM1,2 protein was purified via nickel affinity chromatography on a Hi-trap column (GE Biosciences) followed by a Hi-trap Q column (GE Biosciences). The (His)6 purification tag was cleaved using the tobacco etch virus (TEV) enzyme and the cleavage mixture was then loaded onto Hi-Trap columns (GE Biosciences) to isolate the protein. Subsequently, A-RRM1,2 was loaded onto a HiLoad 16/600 Superdex 75 pg (GE Bioscience) gel filtration column and eluted into the desired buffer. Protein stock solutions were kept in a buffer consisting of 10 mM K2HPO4, 0.5 mM EDTA, 20 mM KCl, and 4 mM BME (pH 6.5).

The UP1 protein (A1-RRM1,2) used in this study (residues 1–196) was subcloned into a pET28a vector and subsequently overexpressed as a C-terminal (His)6-tagged fusion protein in BL21 (DE3) cells and grown in LB Broth. The C-terminal (His)6-UP1 was purified via nickel affinity chromatography on a Hi-Trap column (GE Bioscience) and subsequently loaded onto a HiLoad 16/600 Superdex 75 pg (GE Bioscience) gel filtration column and eluted into the desired buffer. Protein stock solutions were kept in a buffer consisting of 10 mM K2HPO4, 120 mM NaCl, 0.5 mM EDTA, and 5 mM DTT at pH 6.5. Protein homogeneity was confirmed by SDS-PAGE and concentrations were determined using the theoretical molar extinction coefficient.

### NMR Data Acquisition

NMR spectra were recorded on a Bruker Avance (700 and 900 MHz) high-field spectrometer equipped an HCN triple resonance cryoprobe and a z-axis field gradient accessory. All 2D NMR data were processed with nmrPipe/nmrDraw and analyzed using NMRFx analyst. (*23, 24*) Hydrogen-bonding was assigned by collecting exchangeable ^1^H imino spectra in 90% H2O, 10% D2O buffer containing 10 mM K2HPO4 (pH 6.5), 20 mM KCl, 0.5 mM EDTA, and 4 mM TCEP at 283 K using a Watergate NOESY (*t*m = 200 and 250 ms) pulse sequence on the fully protonated RNA constructs.

### Isothermal Titration Calorimetry

RNA and protein samples used for calorimetry were prepared as described above. Calorimetric titrations were performed on a VP-ITC calorimeter (Microcal, LLC) at 25 °C using 10 mM K2HPO4, 20 mM KCl, 0.5 mM EDTA, and 4 mM BME (pH 6.5) buffer centrifuged and degassed under vacuum before use. The respective A1-RRM1,2 at 100 μM was titrated into ∼1.4 mL of the respective RNA construct (SLII, SLII^CCC^, SLII^resist^) at 4 μM over a series of 32 injections set at 6 μL each. To minimize the accumulation of experimental error associated with batch-to-batch variation, titrations were performed in duplicate. Data were analyzed using the KinITC routines supplied with Affinimeter. (*25*)

For the competition experiments, calorimetric titrations were performed on a VP-ITC calorimeter (Microcal, LLC) at 25 °C into 10 mM K2HPO4, 20 mM KCl, 0.5 mM EDTA, and 4 mM BME (pH 6.5) buffer centrifuged and degassed under vacuum before use. A-RRM1,2 at 100 μM was titrated into ∼1.4 mL of 4 μM SLII:UP1 complex at a 1:1.2 molar ratio over a series of 32 injections set at 6 μL each. Likewise, UP1 at 100 μM was titrated into ∼1.4 mL of 4 μM SLII:A-RRM1,2 complex at a 1:3.5 ratio over a series of 42 injections set at 6 μL each. Additionally, this competition experiment was performed in the presence of DMA-135. UP1 at 100 μM was titrated into ∼1.4 mL of 4 μM SLII:A-RRM1,2:DMA-135 at a 1:4:5 ratio, over a series of 42 injections set at 6 μL each. To minimize the accumulation of experimental error associated with batch-to-batch variation, titrations were performed in duplicate. Data were analyzed using KinITC routines supplied with Affinimeter. (*25*)

### Cells and Viruses

SF268 (human glioblastoma) cells were cultured at 37 °C with 5% CO2 in RPMI medium supplemented with 10% FBS (Gibco). Vero (African green monkey kidney) and RD (human embryonal rhabdomyosarcoma) cells were grown at 37 °C with 5% CO2 in MEM medium supplemented with 10% FBS. EV-A71 (TW/2231/98) was propagated in RD cells. EV-D68 (TW-02795-2014) was propagated in Vero cells.

### Isolation of (DMA-135)-resistant Virus, 5’UTR Cloning, and Sequencing

SF268 cells were seeded in 6-well plate at a density of 3 x 10^5^ per well and cultured for 24 hrs prior to virus infection. Cells were infected with undiluted EV-A71 stock (designated passage 0 virus) in the presence of 50 μM DMA-135 in RPMI supplemented with 2.5% FBS for 24 hrs at 37 °C. Medium was collected and saved (passage 1 virus). A 0.5 ml portion of medium was used to infect fresh SF268 cells again for 24 hrs. Medium was harvested (passage 2). This process was repeated to passage 10. Virus titers of successive passages were determined by plaque assay using Vero cells. (*26*) Passages 9 and 10 virus had titers comparable to passage 0 virus. Passage 9 virus was further characterized. Vero cells were infected with passage 9 virus and overlaid with 1% low melting agarose (Invitrogen) in MEM, 2.5 % FBS at 37 °C for 4 days. Plaques were picked, virus was eluted, and virus titers were determined by plaque assay. One plaque-purified virus sample was selected for infection of SF268 cells for 24 hrs. Total RNA was extracted from infected cells using a RNeasy kit (Qiagen). The 5’UTR region of EV-71A RNA was amplified using RT-PCR with primers EV1F: 5’-TTAAAACAGCCTGTGGGTTGC and EV745R: 5’-GTTTGATTGTGTTGAGGGTCA. The amplified cDNA fragment was cloned into plasmid pCRII-TOPO by TA cloning (Life Technologies). The sequence of the 5’UTR cDNA was determined using primer EV1F and compared to the sequence of the infectious clone used to prepare passage 0 virus.

### Construction of EV-A71 mutant with nt132 and nt133 mutations

The infectious clone of EV-A71, plasmid pEV71, and the dual luciferase reporter plasmid with nt 132 and nt 133 mutations, pRHF-EV71-5’UTR nts 132-133, were both digested with Apa I and Msc I. The ApaI-MscI fragment from pRHF-EV71-5’UTR nt132nt133 was purified and ligated into plasmid pEV71 digested with the same restriction enzymes. The resulting colonies were screened by sequencing to identify those harboring the two mutations. To prepare DNA for in vitro transcription, plasmid from a positive colony was purified and digested with EcoR I, fractionated in an agarose gel, and the infectious clone fragment purified. Full-length viral RNA containing the nt 132 and nt 133 mutations was synthesized by in vitro transcription using the infectious cDNA fragment. RNA was purified and transfected into SF268 cells. Supernatant was harvested 3 days after transfection. The mutant virus titer was determined by plaque formation with Vero cells.

To determine the effect of DMA-135 on mutant virus growth, SF268 cells were infected with wild-type or mutant virus at an moi = 1 for 24 hrs ± 50 µM DMA-135. Media were harvested and virus titers were determined by plaque assays with Vero cells.

### Dual Luciferase Reporter Assay

The mutant 5’UTR in pCRII-TOPO was amplified using primers EV1F and EV745R each containing a 5’ NotI restriction site. The PCR fragment was digested with NotI and cloned into the NotI site of pRHF to generate plasmid pRHF-EV71-5UTR-nts 132-133 mut for luciferase assays.(*27*) To assess the effects of putative suppressor mutations on IRES-dependent translation, capped and polyadenylated RNAs were prepared by in vitro transcription from the wild-type and mutant template DNAs linearized using AfeI.(*26*) We refer to these RNAs as RLuc-EV71-5’UTR-FLuc for wild-type and RLuc-EV71-5’UTR^revert^- FLuc for the RNA containing the mutations in SLII. The RNAs were transfected into SF268 cells cultured without (i.e., vehicle) or with 50 μM DMA-135 for 2 days. Dual luciferase assays were then performed using the Dual Luciferase Assay kit (Promega) as described.(*26*)

### Biotin-Streptavidin Pulldown Assay of SLII-Protein Complexes

To assess the effects of putative suppressor mutations and DMA-135 on protein-SLII interactions, RNAs were prepared by in vitro transcription from wild-type and mutant SLII template DNAs in the presence of biotin-labeled UTP. RNAs were capped and polyadenylated to maintain stability of the RNAs in cells. These RNAs were transfected into SF268 cells, and after 4 hrs, various concentrations of DMA-135 were added to culture media. Twelve hrs post-transfection, cell lysates were prepared and SLII-protein complexes were captured using streptavidin-sepharose. AUF1 and hnRNP A1 bound to SLII were detected by Western blot analyses. (*26*)

## Results

### Selection of DMA-135 resistant EV-A71 virus harboring mutations in the SLII IRES domain

As previously reported, DMA-135 functions as an inhibitor of EV-A71 translation and replication. (*15*) We further demonstrated that the small molecule targets the bulge motif in the SLII IRES domain to induce a conformational change to the RNA structure within and adjacent to the bulge loop. Notably, the changes disrupt the A133-U163 base pair and abrogate sequential stacking of the AAU sequence motif of the bulge. The exposed AAU motif forms part of the high-affinity binding site for AUF1. (*15*)

To evaluate the *in vivo* mechanism of action of DMA-135, we grew (DMA-135)-resistant viruses by repeated culturing of the wild-type virus for 10 passages at a fixed concentration (50 μM) of the inhibitor. The virus titers for each passage were determined by plaque assay (**Fig. 1A**). Initial resistance to DMA-135 was observed after passage 4 (3.9𝑥10^’^ 𝑝𝑓𝑢/𝑚𝐿). This resistance improved over passages 5 to 9 (8.4𝑥10^1^ 𝑝𝑓𝑢/𝑚𝐿). Interestingly, the virus titers obtained in passage 9 were comparable to those obtained when using the wild-type EV-A71 strain in the absence of the inhibitor (**Fig. 1A**), indicating the generation of a (DMA-135)-resistant EV-A71 revertant

**Figure 1.**
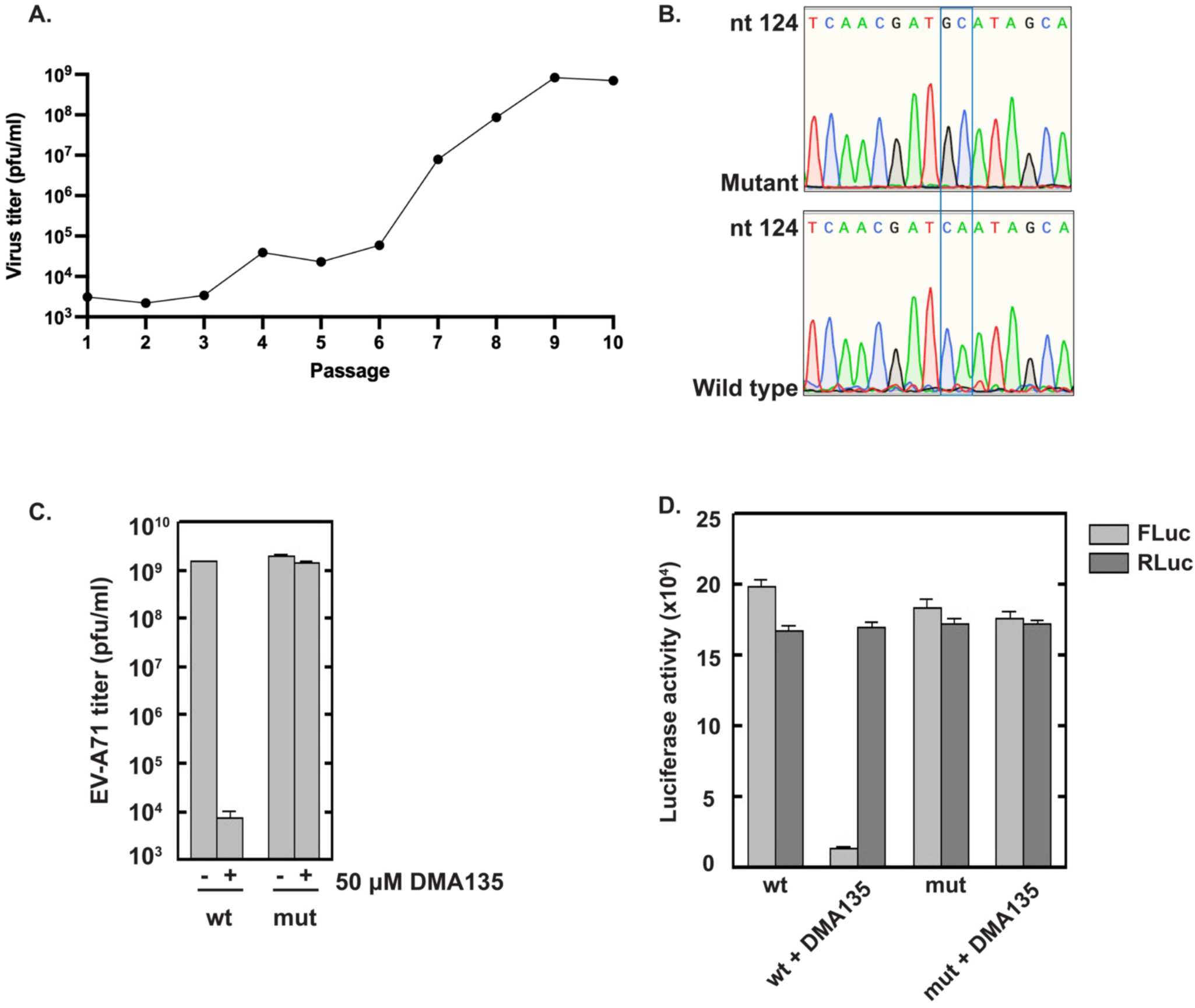
Pressure-induced viral evolution selects for revertant viruses resistant to DMA-135. (**A**) Serial passaging of EV-A71 in the presence of 50 µM DMA-135. (**B**) Sequence comparison of wild-type and the revertant EV-A71 genome selected after serial passage 10. The region shown corresponds to the 5’-half of SLII that includes the bulge loop. The sequence comparison shows that the revertant virus incorporated two nucleotide substitutions (C132G and A133C) in SLII. (C) A mutant EV-A71 virus that incorporates the C132G and A133C mutations only is refractory to DMA-135 at concentrations that robustly inhibit the wild-type virus. (D) EV-A71 IRES with C132G and A133C mutations in SLII retain normal activity levels in the presence of DMA-135 concentrations that inhibit FLuc activity that is driven by the wild-type IRES.

We next characterized the (DMA-135)-resistant EV-A71 mutant obtained at passage 9. Following purification of virus from one selected plaque and RT-PCR of the 5’UTR, the resulting DNA was sequenced in order to identify mutations that may confer its resistance to DMA-135. Figure 1B summarizes the sequencing results. The two mutations identified mapped to the bulge loop region of the SLII domain. Particularly, C132 and A133 were substituted to G132 and C133 in the revertant virus (C132G, A133C). These nucleotides are immediately adjacent to the bulge structure of SLII (hereafter referred to as SLII^resist^; see **Fig. 1B** and **Fig. 2A**). To evaluate if the C132G and A133C mutations alone were sufficient to confer drug resistance, we performed site-directed mutagenesis of the wild-type infectious clone cDNA of EV-A71 to introduce the two nucleotide substitutions. Transfection of Vero cells with the resulting EV-A71 mutant RNA, prepared by *in vitro* transcription, was used to generate mutant virus harboring only the two SLII mutations. Subsequently, SF268 cells were infected with wild-type or mutant virus in parallel, with or without 50 µM DMA-135. Figure 1C shows that the C132G, A133C mutant virus is refractory to DMA-135 at this concentration compared to wild-type EV-A71. These results prove that the bulge loop environment of SLII is the biologically relevant target of DMA-135 and that the mutations identified are sufficient to confer DMA-135 resistance to EV- A71.

**Figure 2.**
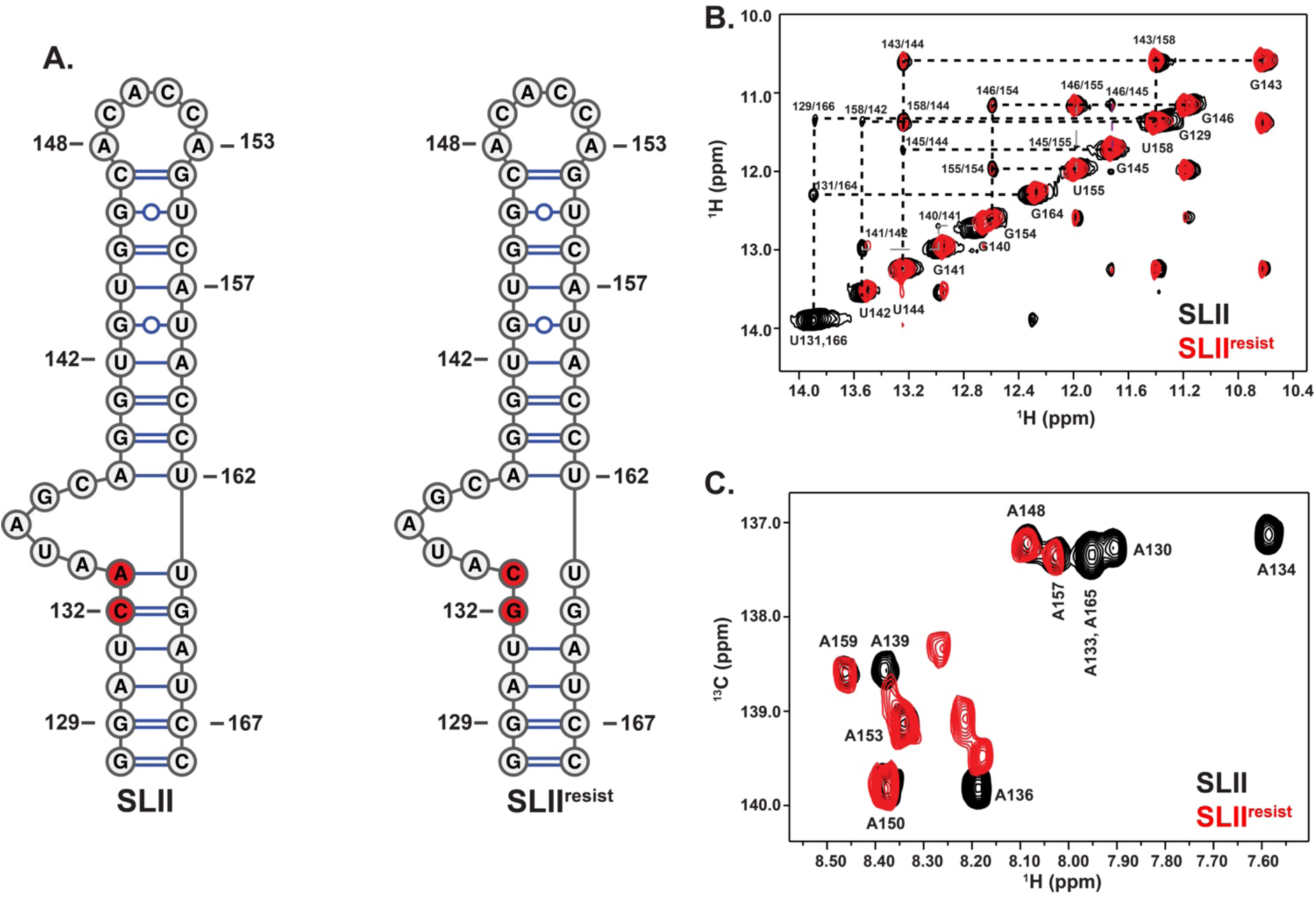
SLII^resist^ folds into a global structure with similar features to that of wild-type SLII. (**A**) Comparison of the experimentally-determined SLII (left) structure to that of SLII^resist^ (right). (**B**) Overlay of the ^1^H-^1^H NOESY spectra of SLII (black) and SLII^resist^ (red) reveals the two RNAs have identical (G/U)NH-(G/U)NH cross peaks patterns for nucleotides corresponding to the apical loop and upper helix. NOE cross peaks corresponding to U131 and U166 located in the lower helix of SLII are missing in SLII^resist^ providing evidence that the C132G and A133C mutations change the local structure near the bulge loop. The spectra were recorded at 900 MHz in 10 mM K2HPO4 (pH 6.5), 20 mM KCl, 0.5 mM EDTA and 4 mM BME D2O buffer at 298 K. (**C**) Overlay of the TROSY HSQC spectra of A(^13^C)-selectively labeled SLII (black) and SLII^resist^ (red) demonstrate that the base stacking arrangements of the two RNAs are similar within the upper apical loop region but differ within the bulge loop. The spectra are centered on the C8-H8 region and were collected in 10 mM K2HPO4 (pH 6.5), 20 mM KCl, 0.5 mM EDTA and 4 mM BME D2O buffer at 298 K.

### DMA-135 no longer inhibits IRES-dependent translation in EV-A71 harboring the SLII^resist^ mutations

Building on the identification of the C132G, A133C resistant mutations in the SLII IRES domain, we next evaluated whether DMA-135 affected IRES-dependent translation using a dual-luciferase reporter assay with lysates of cells transfected with EV-A71 IRES-driven reporter RNAs. Reporter plasmids harboring the wild-type or resistant 5’UTR (5’UTR^resist^) linked to firefly luciferase served as templates for *in vitro* synthesis of capped and polyadenylated reporter RNAs, RLuc-(EV-A71/5’UTR)-FLuc and RLuc-(EV- A71/5’UTR^resist^)-FLuc, respectively (see Materials and Methods). The 5’ ORF in both RNAs is Renilla luciferase (RLuc), the translation of which is cap-dependent and thus serves as an internal control. The respective RNAs were transfected into SF268 cells cultured in the absence or presence of DMA-135 (50 μM). The activities of Renilla (RLuc) and Firefly (FLuc) were measured 2 days post-transfection. Figure 1D summarizes the luciferase assay results. As previously reported, DMA-135 attenuates IRES-dependent translation (FLuc) with no significant effects on cap-dependent translation (RLuc) with the wild-type 5’UTR reporter. Particularly, FLuc activity declined by 94% with 50 μM DMA- 135 while RLuc activity remained constant with or without DMA-135. Conversely, FLuc activity was unaffected by 50 μM DMA-135 with the 5’UTR^resist^-containing reporter RNA. As expected, control RLuc was unaffected by DMA-135 indicating that DMA-135 has no effect on cap-dependent translation. These collective results strongly support that the mechanism by which DMA-135 attenuates IRES-dependent translation is via binding to the bulge loop surface of SLII, as we showed previously by NMR spectroscopy. (*15*)

### Resistant mutations alter the structure of the SLII bulge but not its apical loop

To better understand how the C132G and A133C mutations in SLII contribute to DMA- 135 resistance, we proceeded to study the structural properties of this RNA construct by NMR spectroscopy. We confirmed the secondary structure of SLII^resist^ (**Fig. 2A**) by comparing its ^1^H-^1^H NOESY to that of the wild-type construct. Figure 2B shows that identical NOE cross peaks are observed for the upper helices of both wild-type SLII and SLII^resist^ indicating that the C132G and A133C mutations do not perturb the apical loop environment. By contrast, the sequential NOEs involving the U131/U166 spin systems seen in the spectrum of wild-type SLII are completely missing in SLII^resist^, verifying that the revertant mutations change the secondary structure of the bulge proximal region.

Further evidence of localized structural differences in SLII^resist^ relative to wild-type was verified by comparing ^1^H-^13^C HSQC spectra collected on samples prepared with A(^13^C)- selective labeling. Figure 2C shows an overlay of the C8-H8 region of the ^1^H-^13^C TROSY HSQC of SLII^resist^ and SLII. The C8-H8 correlation signals belonging to the adenosines of the upper helix (A157 and A159) and the apical loop (A148, A150 and A153) in the SLII^resist^ construct overlay perfectly with those observed in wild-type SLII. The correspondence between the ^1^H and ^13^C chemical shifts of these residues indicates that the C132G and A133C mutations do not perturb base stacking interactions within and nearby the apical loop. By contrast, the NMR signals of A130, A134, A136, A139 and A165 - bases within or proximal to the bulge - shift to new positions within the spectrum (note A133 becomes C133 in SLII^resist^). The different chemical shifts of these residues relative to the wild-type SLII construct is evidence that the mutations change the local physicochemical environment of the bulge. Collectively, the NMR results reveal that viral evolution, driven by the selective pressure of DMA-135, leads to a variant with structurally perturbing mutations localized to the bulge loop environment of SLII^resist^.

### DMA-135 binds SLII^resist^ but it is unable to induce the formation of a ternary complex

Since the C132G and A133C mutations change the local bulge loop structure, we evaluated if DMA-135 retains binding affinity for SLII^resist^. Using the A(^13^C)-selectively labeled SLII^resist^ construct, we performed a single-point ^1^H-^13^C TROSY HSQC titration as described previously. (*15, 28*) Given the labeling strategy, we are able to assess the extent to which DMA-135 interacts with specific surfaces on SLII^resist^ because the degree of NMR signal perturbations is a proxy for complex formation. (*15, 28*) Figure 3A shows the effects of the addition of excess (5:1) DMA-135 on the C8-H8 correlation signals of SLII^resist^. The spectrum reveals that several of the correlation peaks exchange broaden in the presence of excess DMA-135; however, the signals that overlap with nucleotides from the apical loop and upper helix of wild-type SLII remain mostly unperturbed (A148, A150, A153 and A157). This observation indicates that although the local structure of the bulge motif has changed, DMA-135 can still recognize it as a binding surface.

**Figure 3.**
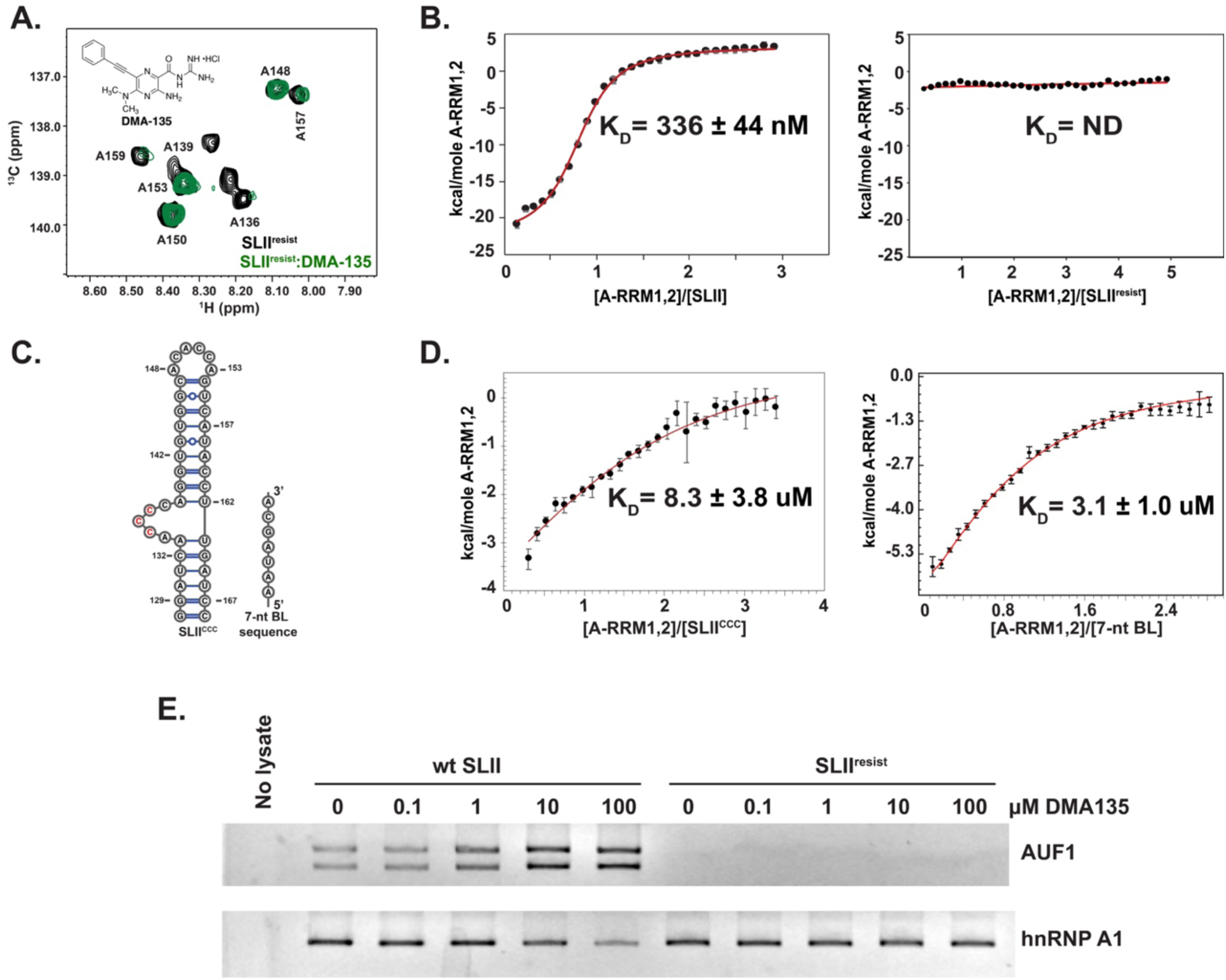
The sequence and structure of the SLII bulge loop facilitates intermolecular interactions. (A) Single-point TROSY HSQC titration of A(^13^C)-selectively labeled SLII^resist^ with DMA-135 at a 5-fold excess. The black correlation peaks correspond to free SLII^resist^ and the red to the (DMA-135)-SLII^resist^ complex. The spectra were collected at 900 MHz in 10 mM K2HPO4 (pH 6.5), 20 mM KCl, 0.5 mM EDTA and 4 mM BME D2O buffer at 298 K. (B) Calorimetric titrations of A-RRM1,2 into SLII (left) and SLII^resist^ (right). The A-RRM1,2-SLII data were fit to a 1:1 stoichiometric binding model in Affinimeter. (*25*) Reported values for dissociation constants (KD) and corresponding standard deviation are from triplicate experiments. (C) RNA constructs used to assess determinants of specific and high-affinity AUF1-SLII interactions. Left, SLII^CCC^ replaces the phylogenetically conserved UAG bulge motif with CCC. Right, a 7-nt oligonucleotide that mimics the SLII bulge loop sequence with adjacent nucleotides. (D) Calorimetric titrations of A-RRM1,2 into SLII^CCC^ (left) and the 7-nt oligonucleotide (right). The A-RRM1,2-SLII^CCC^ data were fit to a 1:1 stoichiometric binding model in Affinimeter. (*25*) Reported values for dissociation constants (KD) and corresponding standard deviation are from triplicate experiments. (E) Protein-biotinylated RNA pull-down experiments were performed to evaluate the influence of DMA-135 on the interaction between AUF1 and SLII^resist^. Biotinylated SLII and SLII^resist^ were transfected into SF268 cells. Cells were cultured with increasing concentrations of DMA-135. Cell lysates were used for pull-down assays of SLII-associated proteins and detected by western blotting.

Because DMA-135 can still interact with the bulge loop environment, we next decided to test if it can allosterically increase the affinity of AUF1 for SLII^resist^ like it does for the wild-type SLII. (*15*) We posited that the mechanism of action (functional specificity) of DMA- 135 is its ability to shift the AUF1-SLII equilibrium to favor suppression of IRES-dependent translation. Figure 3B shows a calorimetric titration of the RNA binding domain of AUF1 (A-RRM1,2) titrated into wild-type SLII and SLII^resist^. As previously observed, the thermogram shows that A-RRM1,2 binds SLII as a specific 1:1 complex with high affinity (KD = 336±44 nM). By contrast, the thermogram of A-RRM1,2 titrated into SLII^resist^ is flat as indicated by no change in the total binding enthalpy over the course of the titration. This observation implies that C132 and/or A133 are determinants of high-affinity A1- RRM1,2-SLII recognition.

To investigate the sequence and structure contributions further, we performed calorimetric titrations of A-RRM1,2 with a lab-derived SLII mutation where the central UAG motif is mutated to CCC (SLII^CCC^) and a synthetic 7-nt oligo (5’-AAUAGCA-3’) that mimics the native bulge sequence and adjacent A133 and A139 nucleotides (**Fig. 3C**). Figure 3D shows that A-RRM1,2 binds SLII^CCC^ very weakly (KD=8±3 µM) and non-specifically as determined by the shape of the binding isotherm relative to the wild-type SLII control titration. This observation is consistent with prior results where we were unable to detect AUF1 in pulldowns using a biotinylated SLII^CCC^ RNA. (*15*) By comparison, the binding affinity of A-RRM1,2 for the 7-nt oligo is also weak (KD=3±1 µM) but slightly tighter than that determined for SL^CCC^. When interpreted collectively, the titrations of A-RRM1,2 with wild-type SLII, SLII^resist^, SLII^ccc^ and the 7-nt oligo reveal the importance of the bulge loop structure, its sequence, and the contributions of the A133-A134 dinucleotide to high-affinity binding.

We decided to test if the physicochemical property of SLII^resist^ to no longer bind AUF1 productively also occurred in a more biological context using an assay where SLII^resist^ was biotinylated and used to pull-down endogenous AUF1. Biotinylated SLII^resist^ and wild-type SLII RNAs were transfected into SF268 cells in the presence of various concentrations of DMA-135. Twelve hours post-transfection, cells were lysed, and the complexes of cellular proteins bound to SLII^resist^ and SLII wild-type were captured using streptavidin-sepharose conjugated beads; levels of captured AUF1 were assessed by Western blot; hnRNP A1 served as a control. Consistent with the calorimetric results, we were unable to detect a complex between SLII^resist^ and endogenous AUF1 whereas the AUF1-SLII wild-type complex readily increased with DMA-135 concentration as we showed previously (**Fig. 3E**). (*15*) By contrast, binding by hnRNP A1 to SLII^resist^ was readily detected with both the wild-type and revertant SLII (also see next section below), though there was a slight decrease in binding by hnRNP A1 to native SLII at the highest DMA-135 concentration tested, 100 µM (**Fig. 3E**). These results solidify the allosteric mechanism of DMA-135 and it shows that EV-A71 can evolve to shift the binding affinity of AUF1 for SLII when under selective pressure.

### DMA-135 modulates the SLII regulatory axis to abrogate robust hnRNP A1 interactions

Stem loop II functions as a regulatory axis during EV-A71 replication by modulating the efficiency of IRES-dependent translation. Several ITAFs and a viral derived small RNA, vsRNA1, bind to SLII to differentially control IRES activity, and DMA-135 overrides these regulatory mechanisms to repress viral translation. (*11, 15–21*) To better understand how the physicochemical properties of SLII contribute to EV-A71 biology, we carried out a series of calorimetric titrations using the RNA binding domains of hnRNP A1 (UP1) and AUF1 (A-RRM1,2). Titrations of UP1 into SLII^resist^ resulted in a biphasic isotherm similar to that previously observed for wild-type SLII (**Fig. 4B**). Fitting of the processed data to a two-independent site binding model reveals that both events are characterized by nanomolar affinities (KD1=2.4±0.4 nM, KD2=247±38 nM), which are on the same order of magnitude as those measured here for wild-type SLII (KD1=0.5±0.1 nM, KD2=178.1±22.7 nM). This result is consistent with the data described above where hnRNP A1 retains binding affinity for SLII^resist^ (±DMA-135) within a biological context (**Fig. 3E**). These collective observations indicate that the C132G and A133C mutations do not significantly impair the hnRNP A1-SLII arm of the regulatory axis, allowing hnRNP A1 to continue to stimulate translation during the earliest stages of viral replication.

**Figure 4.**
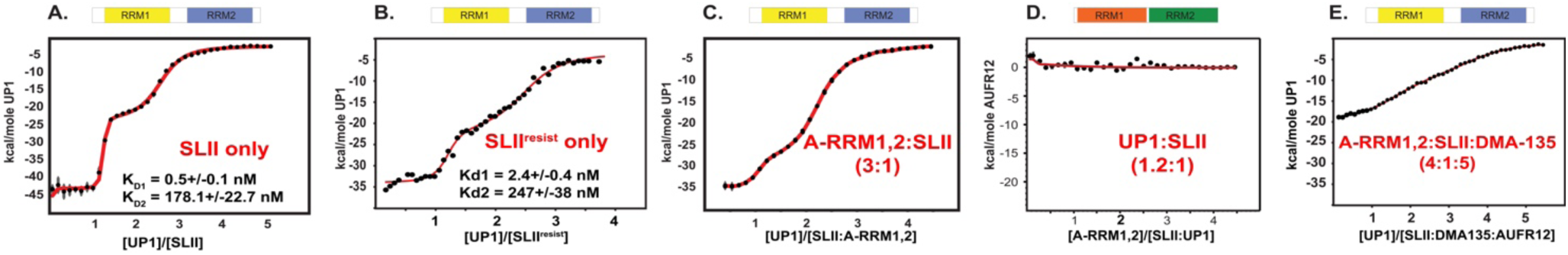
AUF1 and hnRNP A1 compete for the bulge loop surface on SLII. (A-E) Comparative calorimetric titrations of the RNA binding domains of hnRNP A1 (UP1 - yellow/blue rendering) and AUF1 (A-RRM1,2 – orange/green rendering) demonstrate that the two proteins directly compete for the bulge loop surface on SLII and that the degree of the competition is modulated by DMA-135. Thermodynamic parameters derived from fits to a two-independent sets of site model using Affinimeter (*25*) are reported for the UP1-SLII and UP1-SLII^resist^ titration only. Thermodynamic parameters are not reported for the other titrations given the complexity of the competitive binding equilibria. Qualitative comparisons demonstrate that the thermodynamic parameters are observably different for UP1 titrated into preformed complexes of AUF1-SLII and AUFI-SLII-(DMA-135) as well as AUF1 titrated into a preformed complex of UP1-SLII.

Given that hnRNP A1, but not AUF1, can still bind SLII^resist^, we reasoned that DMA-135 modulates the SLII regulatory axis by affecting the extent to which these two proteins compete for the bulge loop. To that end, we performed competitive calorimetric titrations of A-RRM1,2 and UP1 into SLII alone or pre-bound in various complexes (**Fig. 4**). As described above, A-RRM1,2 binds SLII as a specific 1:1 complex with a KD∼330 nM; however, binding is undetectable when it is titrated into a preformed UP1:SLII (1.2:1) complex (**Fig. 4D**). By comparison, UP1 can still bind SLII when in the presence of excess A-RRM1,2, albeit with a characteristically different biphasic isotherm compared to the control titration (**Fig. 4A** and **4C**). The initial transition that corresponds to UP1 binding the bulge loop has a shallower inflection and a lower total change in binding enthalpy (**Fig. 4A** and **4C**). Conversely, the portion of the isotherm that reflects binding of UP1 to the apical loop is primarily unchanged. These results show that hnRNP A1 and AUF1 directly compete for the bulge loop surface and that hnRNP A1 can displace AUF1, which is consistent with the more than 500-fold difference in their binding affinities for the bulge loop.

Next, we decided to see if DMA-135 changes the binding properties by titrating UP1 into a preformed (DMA-135-SLII-(A-RRM1,2) ternary complex. Figure 4E shows that UP1 no longer efficiently displaces A-RRM1,2 from the bulge as determined by the shape of the isotherm and change in total binding enthalpy relative to the binary (A-RRM1,2-SLII) titration. These collective results agree with the antiviral mechanism of action of DMA-135, which is to decrease cap-independent translation by stabilizing the repressive AUF1-SLII complex. DMA-135 shifts the (hnRNP A1/AUF1)-SLII regulatory axis just enough such that viral protein synthesis becomes inhibited during the EV-A71 replication cycle.

### DMA-135 is less efficient at inhibiting EV-D68 replication

To better understand the functional specificity of DMA-135, we decided to test if it can also inhibit the related EV-D68. We selected the Fermon strain of EV-D68 for this study, which has a predicted SLII structure (SLII^Fermon^) consisting of an internal loop instead of a bulge, and a structurally similar apical loop to that of EV-A71 SLII (**Fig. 5A**). We confirmed the base pair composition of the SLII^Fermon^ structure by 2D NMR spectroscopy (**Fig. 5B**). Sequential and long-range (G/U)NH-(G/U)NH NOE cross peaks can be traced for the internal base pairs of the lower and upper helices verifying that SLII^Fermon^ adopts an overall topology consistent with the predicted structure (**Fig. 5B**). Figure 5C shows that addition of 100 µM of DMA-135 to Vero cells infected with EV-D68 (moi=1) reduced viral titers by approximately four orders of magnitude relative to the vehicle (DMSO) control (**Fig. 5C, left**). This suggests that the slight differences in the SLII structures (**Fig. 5A**) account for the lower efficacy of DMA-135 for EV-D68 relative to EV-A71, which decreased almost six orders of magnitude at 50 µM DMA-135 compared to the vehicle control (**Fig. 5C, right**). As a further probe of the structural differences, we performed calorimetric titrations of A-RRM1,2 into SLII^Fermon^. The sequence composition of the 5’-half (putative location of the AUF1 binding site) of the internal loop that aligns with the bulge loop (AAUAGCA) of EV-A71 SLII is UUAGAA. Figure 5D shows that A-RRM1,2 binds to SLII from EV-D68 with significantly weaker affinity (KD=846±72 nM) compared to EV-A71. Notably, DMA-135 has a modest effect on the binding affinity (KD=556±62 nM) when A-RRM1,2 is titrated into a preformed (DMA-135)-SLII^Fermon^ complex. Thus, we conclude that the binding of DMA-135 to EV-A71 SLII is highly specific and that the physicochemical properties of EV-A71 SLII determine the functional specificity of DMA-135.

**Figure 5.**
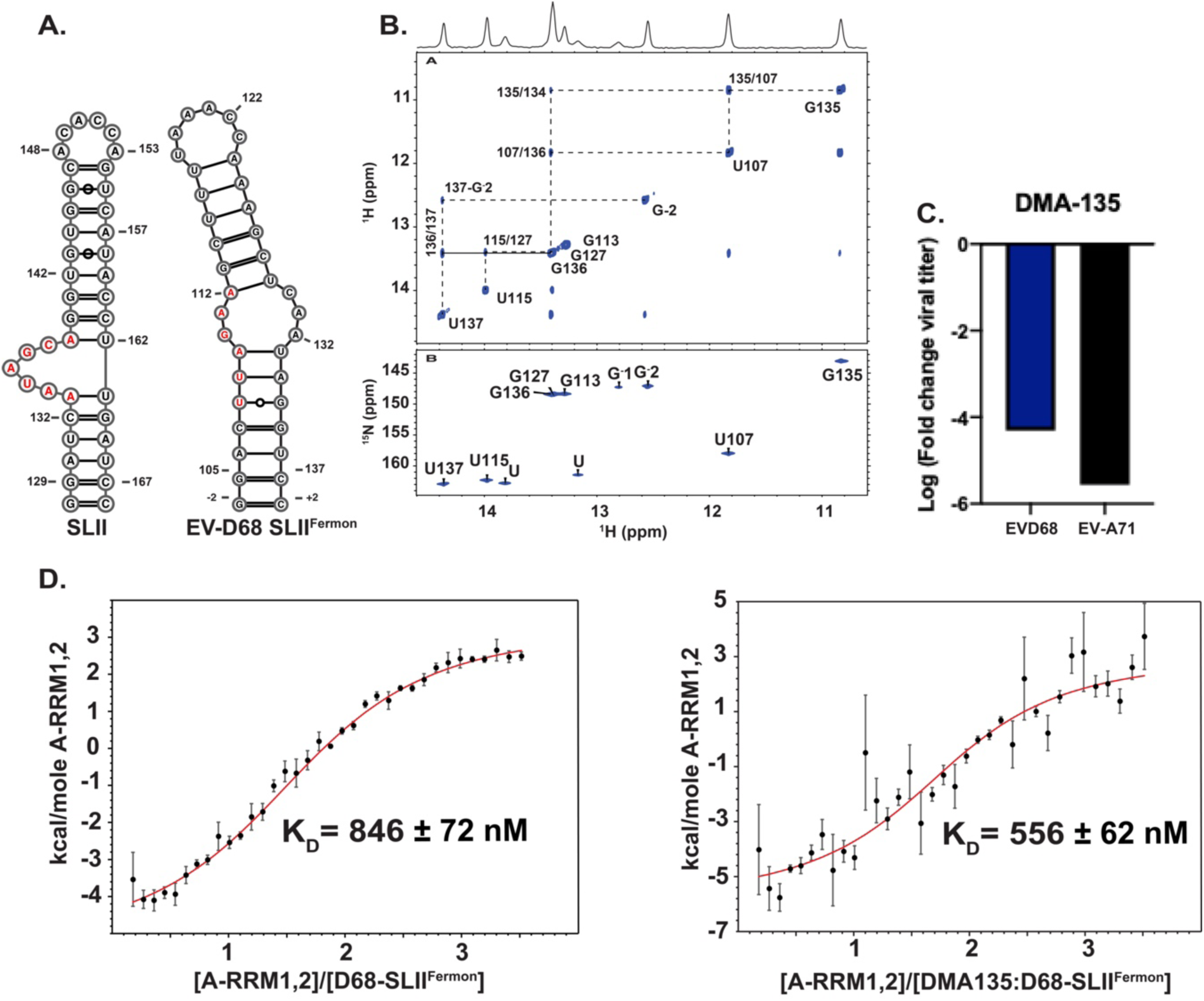
Assessment of the antiviral capacity of DMA-135 against EV-D68. (A) Comparison of the experimental secondary structure of SLII from EV-A71 (left) to that predicted for EV-D68 (right). (B) Top, imino region of the ^1^H-^1^H NOESY (700 MHz and tm=200 ms) spectrum of EV-D68 SLII collected in 10 mM K2HPO4 (pH 6.5), 20 mM KCl, 0.5 mM EDTA, and 4 mM TCEP with 10% D2O, at 283K. The vertical and horizontal dashed lines trace the NOE stacking pattern for each stable helical region demonstrating that EV-D68 SLII^Fermon^ is an independently folded domain. Bottom, ^1^H-^15^N HSQC collected in the buffer condition of 10 mM K2HPO4 (pH 6.5), 20 mM KCl, 0.5 mM EDTA, and 4 mM TCEP with 10% D2O, at 283K, confirms the^1^H-^1^H NOESY assignments. (C) Comparison of the antiviral activity of DMA-135 against EV-A71 (50 µM DMA-135) and EV-D68 (100 µM DMA-135). Log10 values of fold change in virus titers compared to the DMSO (vehicle) control are shown. (D) Calorimetric titrations of A-RRM1,2 into (left) EV-D68 SLII^Fermon^ and (right) a preformed (1:5) SLII^Fermon^-(DMA-135) complex. The titration data were fit to a 1:1 stoichiometric binding model in Affinimeter. (*25*) Reported values for dissociation constants (KD) and corresponding standard deviation are from triplicate experiments.

## Discussion

Viral evolution under selective pressures of small molecules with capacity to inhibit replication offers opportunities to elucidate mechanisms of action and to reveal new biology. For small molecules that target viral RNA structures, pressure-driven evolution can also inform on determinants of functional specificity even within a background of potentially non-productive binding events. Thus, viruses are unique model systems to calibrate principles of small molecule-RNA interactions and to understand the contributions of RNA structures to biological function. Here, we harnessed the power of viral evolution to reveal the cellular mechanism of the antiviral DMA-135 and to characterize host-virus complexes that differentially regulate EV-A71 translation.

Positive strand RNA viruses, like EV-A71 and EV-D68, use the same genomic template for translation and viral RNA synthesis. (*8, 11, 13, 22*) Following infection, translation proceeds in the cytoplasm via internal loading of the ribosome onto the IRES. The efficiency by which ribosomes are recruited to the IRES determines the frequency by which the viral life cycle transitions from early to late stages because viral protein products are necessary to synthesize nascent genomic RNA and to form infectious virions. Nevertheless, high levels of viral proteins are cytotoxic to the cell so positive strand RNA viruses resolve this intrinsic dichotomy by differentially regulating IRES levels via multiple and redundant mechanisms.

Although the complete contributions of the IRES structures of EV-A71 are unknown, the SLII domain is a pivotal regulatory element that coordinates multiple protein-RNA and RNA-RNA interactions to fine-tune viral polyprotein synthesis. (*11, 15–21*) Two such interactions include the recruitment of the host RNA binding proteins, hnRNP A1 and AUF1. These proteins compete for SLII to either stimulate (hnRNP A1) or repress (AUF1) IRES-dependent translation. Genetically introduced mutations that abrogate the SLII-protein regulatory axis robustly restrict EV-A71 replication by attenuating IRES activity, demonstrating the significance of conserved nucleotide epitopes to viral fitness. (*16, 21*)

The observation that DMA-135 resistance was conferred via non-compensatory mutations of C132G and A133C (part of the AUF1 binding site) suggests that the virus naturally exerts less pressure on repressing translation as opposed to stimulating it. Indeed, hnRNP A1 (stimulator) retains binding affinity for SLII^resist^ but AUF1 (repressor) is unable to form a detectable complex with the revertant RNA. As such, EV-A71 escaped sensitivity to DMA-135 by evolving revertant mutations that change the repressive AUF1 arm of the SLII regulatory axis. Furthermore, the activity of the IRES containing SLII^resist^ was comparable to that of the wild-type IRES with or without DMA-135, providing further evidence that inhibition proceeds via selective modulation of the repressive AUF1-SLII complex.

Under non-(DMA-135) conditions, hnRNP A1 alone is able to displace AUF1 pre-bound to SLII; however, it cannot efficiently displace AUF1 when AUF1 is part of a ternary complex with SLII and DMA-135 (**Fig. 4E**). By comparison, AUF1 cannot displace hnRNP A1 when it is already bound to SLII (**Fig. 4D**). These observations imply that the SLII regulatory axis is under thermodynamic control, and mechanisms that modulate the relative SLII-protein binding affinities tune the IRES activity (**Fig. 6**). In support of this concept, DMA-135 dose-dependently decreases IRES-dependent translation by increasing the affinity of AUF1 for SLII. (*15*) Even though DMA-135 retains affinity for SLII^resist^, it can no longer functionally inhibit the revertant virus because the A133C mutation changes part of the AUF1 binding epitope to in turn reduce its binding capacity. When interpreted collectively, the functional mechanism of DMA-135, as validated through viral evolution, is to shift the regulatory axis towards the IRES-repressive SLII-AUF1 arm.

**Figure 6.**
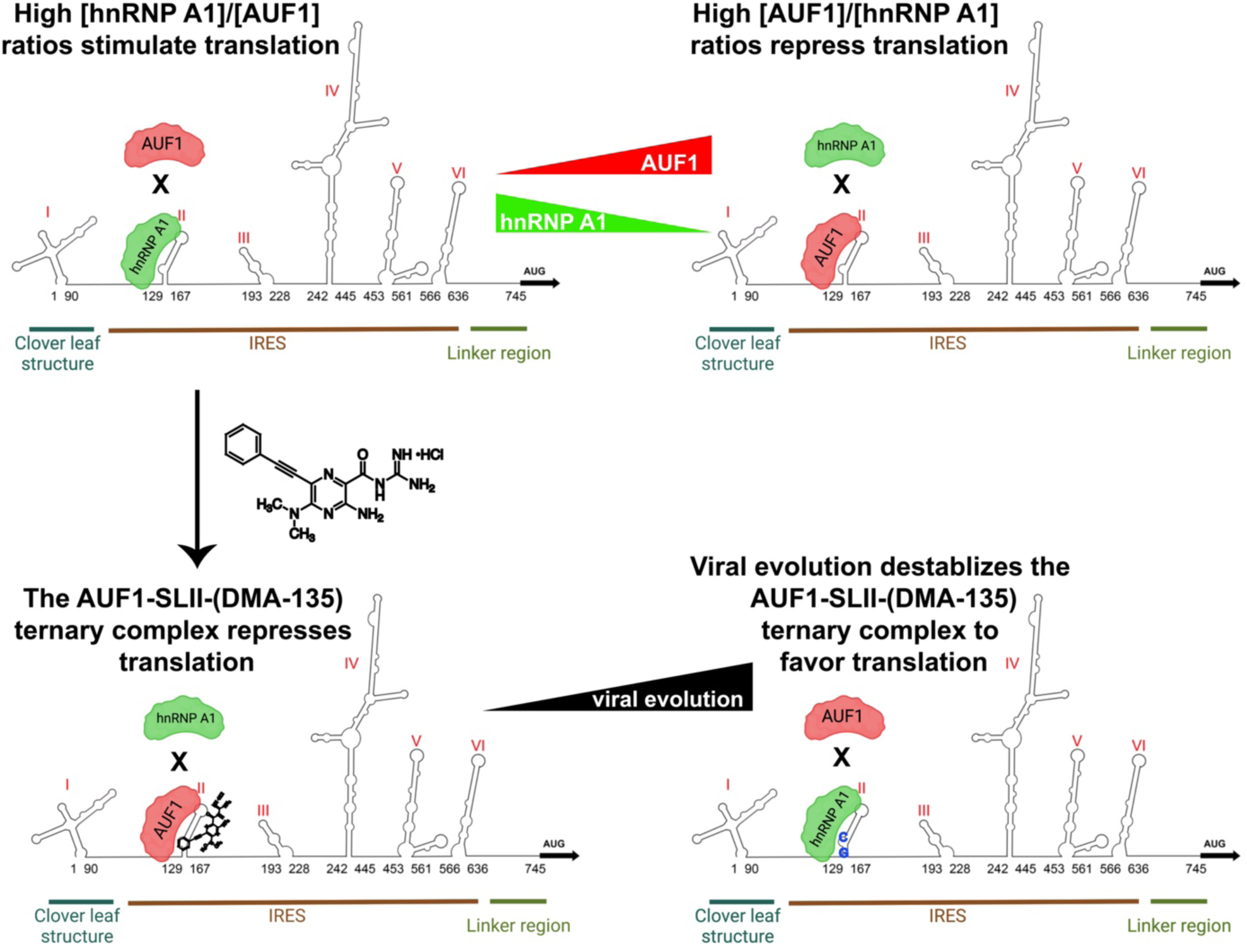
The EV-A71 SLII-hnRNP regulatory axis is under thermodynamic control. An illustration of a conceptual model to interpret the different mechanisms by which the SLII IRES domain coordinates a sub-set of molecular interactions to in turn modulate translational output and viral replication efficiency. Top, under normal conditions of viral infection, hnRNP A1 and AUF1 compete for the bulge loop of SLII to stimulate or repress viral replication, respectively. Factors that change the relative abundance of hnRNP A1 or AUF1 can shift the SLII regulatory axis towards a repressive or stimulatory direction. Bottom, DMA-135 shifts the SLII-hnRNP regulatory axis by stabilizing a ternary complex with AUF1 and SLII to in turn reduce hnRNP A1’s binding capacity. Viral evolution selects for mutations in the AUF1 binding surface on SLII to shift the regulatory axis back towards favoring translation.

We also demonstrated that DMA-135 can inhibit the related EV-D68 virus, albeit ∼100-fold less efficiently compared to EV-A71. Comparison of the structural and biophysical properties of the SLII domains from the two viruses offers insights into the relative differences in DMA-135 efficacy. The SLII domain from EV-A71 consists of a conserved 5-nt bulge loop with adjacent AU base pairs. The sequence of the loop environment is 5’-AAUAGCA-3’, which matches the consensus motifs for AUF1 and hnRNP A1. (*29–31*) By comparison, SLII from the EV-D68 Fermon strain studied here includes an internal loop with a 5’-UUAGAA-3’ motif that best aligns with the sequence of EV-A71. We showed that AUF1 can bind SLII^Fermon^ as a specific and 1:1 complex, albeit ∼3-fold weaker than it forms a complex with SLII from EV-A71. Furthermore, DMA-135 has a modest, yet measurable, influence on the binding affinity of the AUF1-SLII^Fermon^ complex. Together, the data indicate that the potency of DMA-135 against EV-A71 relates to its ability to allosterically recruit AUF1 to an optimal 5’-AAUAGCA-3 bulge loop sequence.

In sum, the work presented here reveals the power of viral evolution to further elucidate functional mechanisms of small molecules with therapeutic capacity against RNA structures. To the best of our knowledge, this is the first example where an RNA-targeting small molecule induced pressure-driven evolution to select for revertant viruses with resistant mutations mapped to the RNA binding site. The work also demonstrates how small molecules can be deployed as chemical biology reagents to interrogate protein-RNA biological interfaces. We believe such tools will have a broad range of utilities to better understand protein-RNA networks, including those found in phase separated compartments.

## Acknowledgements

The authors would like to thank staff at the Northeast Ohio High Field NMR facility for assistance with setting up the experiments. The authors also acknowledge Martina Zafferani for synthesizing DMA-135. Funding was provided by NIH grants R35 GM124785 (AEH, Duke University) and R01 GM126833 (BST, CWRU and GB, Rutgers).

