## Supplemental Information for "Enterovirus Evolution Reveals the Mechanism of an RNA-Targeted Antiviral and Determinants of Viral Replication"

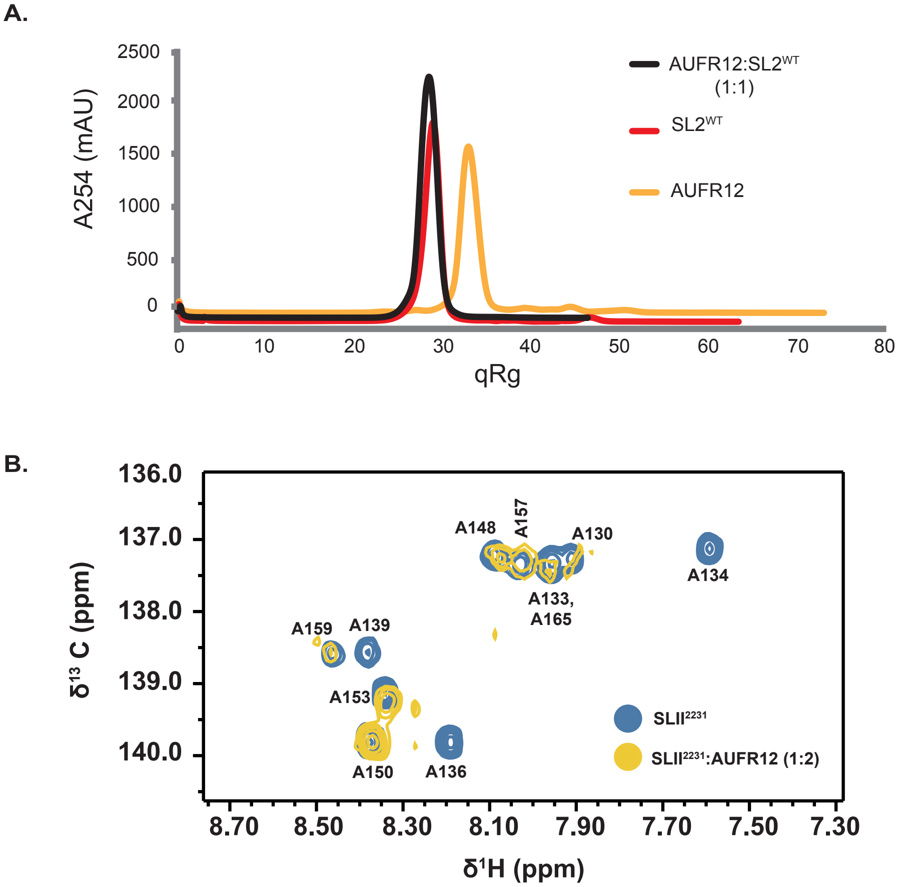


**Figure S1.** Biochemical evidence that A-RRM1,2 forms a stable and robust complex with SLII. (A) Analytical SEC titration of A-RRM1,2 into SLII. Titration was performed by pre-incubating 5 uM of SLII with equal molar amount of A-RRM1,2. The protein-RNA complex was resolved on a Superdex 200 10/300 GL column. (B) $^{1}H$-$^{13}C$ TROSY HSQC titration of A($^{13}C$)-selectively labeled SLII with unlabeled A-RRM1,2. The blue correlation peaks correspond to free SLII and the yellow to the (A-RRM1,2)-SLII complex. The spectra were recorded at 900 MHz in 10 mM $K_2HPO^4$ (pH 6.5), 20 mM KCl, 0.5 mM EDTA and 4 mM BME.

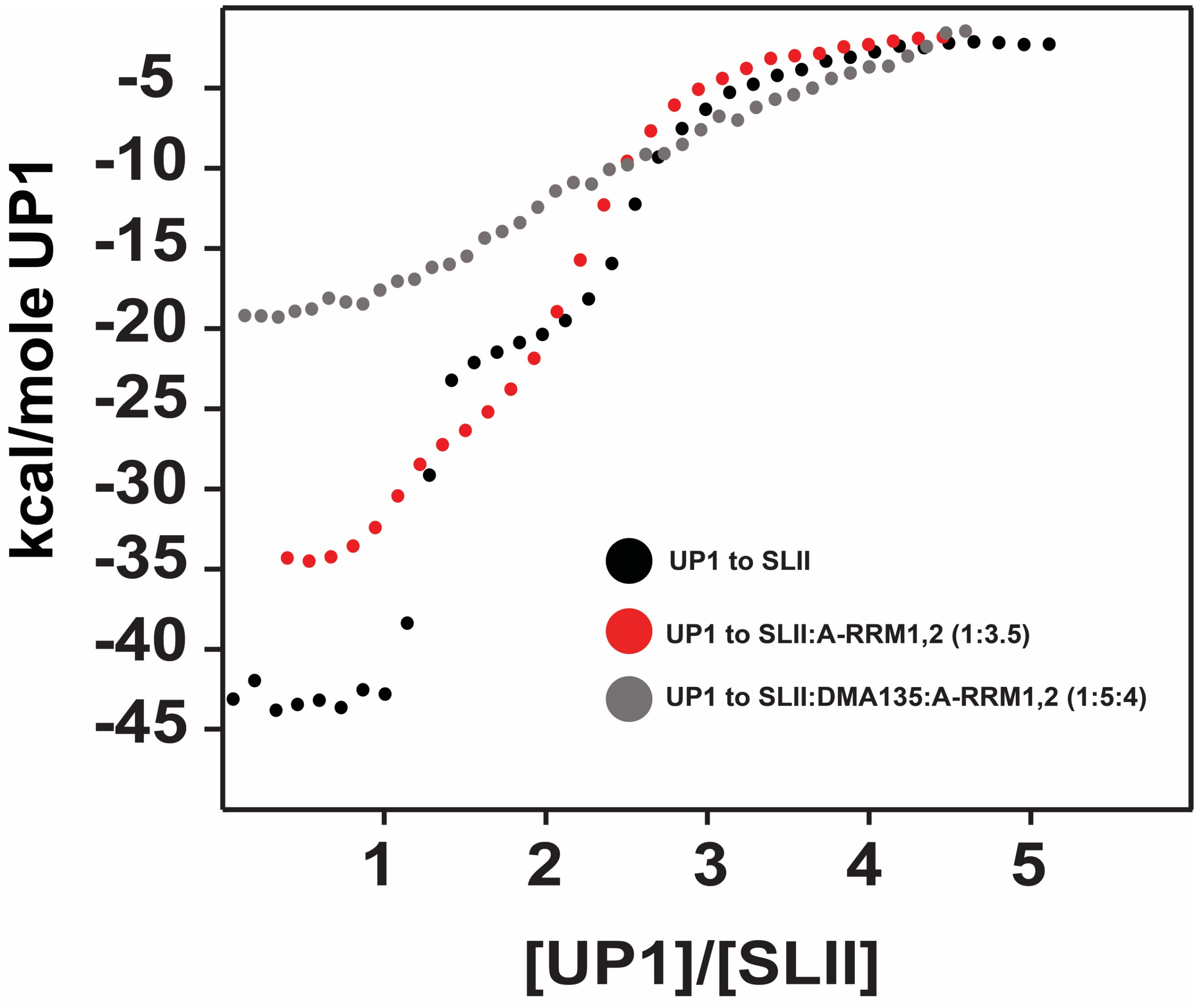


**Figure S2.** Comparison of the calorimetric titrations of UP1 into various complexes of SLII demonstrates competition for the bulge loop environment and the influence of DMA-135 to abrogate UP1 binding productively to the ternary complex. Titrations were performed in 10 mM $K_2HPO^4$ (pH 6.5), 20 mM KCl, 0.5 mM EDTA and 4 mM BME, and at 298K.
